# Nonconscious processing of fearful and neutral faces modulates the N170

**DOI:** 10.1101/2025.01.30.635452

**Authors:** Maximilian Bruchmann, Sebastian Schindler, Pia Breitwieser, Lynn Tilly, Jens Bölte, Torge Dellert, Thomas Straube

## Abstract

Prioritized processing of fearful compared to neutral faces is reflected in differential event-related potentials (ERPs). There is an ongoing debate about the extent to which faces in general and fearful faces in particular enhance specific ERPs if they are not consciously perceived. Specifically, the N170 has been suggested as the most likely candidate for enhanced processing of nonconscious fearful faces. In this pre-registered study, we tested in a large sample based on sequential Bayesian sampling (*N* = 64) whether early components of the ERP (P1, N170, and early posterior negativity; EPN) discriminate between fearful faces, neutral faces, and non-facial control stimuli. Consciousness was manipulated by presenting stimuli for 17 ms in a backward masking design, with the mask following immediately or after a delay of 200 ms. Participants rated their subjective perception on a perceptual awareness scale in each trial. Importantly, only trials in which nothing but the mask was perceived were considered for the analysis of nonconscious effects. The results showed strong evidence for an increased N170 in response to nonconscious fearful compared to neutral faces; however, this difference was significantly smaller than in the conscious condition. Furthermore, we obtained strong evidence for N170 differences between nonconscious faces and no-faces. For P1 and EPN amplitudes, no significant effects were observed in the nonconscious conditions, although exploratory analyses of the P1 peak interval suggest nonconscious face-no-face differentiation. These results support the notion that nonconscious emotion and face processing are detectable in early neural responses and show that these nonconscious effects are considerably weaker than the corresponding conscious effects.

## 1. Introduction

Detecting and responding to emotional facial expressions is highly relevant for survival, especially when these expressions signal a potential threat. In line with this notion, prioritized processing of fearful compared to neutral faces is reflected in behavioral and neural responses, such as event-related potentials (ERPs; e.g., Eimer & Holmes, 2002; Langeslag & van Strien, 2018; Wieser & Keil, 2014). In electrophysiological studies, this prioritization is indexed by the enhancement of several ERP components in response to fearful compared to neutral faces, such as the P1, N170, and early posterior negativity (EPN; for review, see Schindler & Bublatzky, 2020).

The P1 is a positive occipital deflection (peaking between 80 and 100 ms) that is modulated by low-level stimulus features such as contrast (Bobak et al., 1987; Vassilev et al., 1994) or size (Bayer et al., 2012; Busch et al., 2004; Schindler et al., 2018). Regarding face and emotion effects, P1 modulations for faces relative to other stimuli are mixed (Itier & Taylor, 2004; Neumann et al., 2011; but see Ganis et al., 2012; Schindler, Tirloni, et al., 2021). Likewise, increased P1 amplitudes for fearful versus neutral faces are also variable (for a review, see Schindler & Bublatzky, 2020). Both effects are strongly influenced by low-level visual information (Bruchmann et al., 2020; Rossion & Caharel, 2011; Schindler, Bruchmann, et al., 2021). In contrast, the N170, a negative occipito-temporal deflection (peaking at approximately 170 ms), is reliably enhanced for fearful versus neutral faces (Bruchmann et al., 2023; Eimer & Holmes, 2007; Hinojosa et al., 2015; Schindler, Bruchmann, et al., 2021; Schindler, Tirloni, et al., 2021; Schindler & Bublatzky, 2020). The N170 indexes early structural and configural face encoding, and increased N170 amplitudes for faces relative to non-faces have been frequently reported (e.g., Bentin et al., 1996; Itier & Taylor, 2004; Rousselet et al., 2004), even for weak, task-irrelevant stimuli (Dellert et al., 2021; Shafto & Pitts, 2015). Furthermore, fearful faces have been shown to enhance the N170 independent of various physical stimulus manipulations (Bruchmann et al., 2020; Schindler et al., 2019) across a variety of tasks (Mühlberger et al., 2009; E. Smith et al., 2013; Walentowska & Wronka, 2012; Wieser et al., 2012), in the absence of task-relevance of the faces (Schindler, Bruchmann, et al., 2020) and independent of the level of attentional load (Schindler, Caldarone, et al., 2020). Taken together, this suggests that N170 effects of fearful faces reflect highly automatized threat-processing. The EPN typically peaks between 200 and 300 ms and numerous studies have shown it to be increased for fearful compared to neutral expressions (Durston and Itier, 2021; Schindler et al., 2019; Walentowska and Wronka, 2012; Wieser et al., 2012), albeit modulated by attentional tasks (Schindler, Bruchmann, et al., 2020). The EPN has been linked to early attentional selection (Schupp et al., 2004).

It has been suggested that threat information is processed highly automatically (Mineka & Öhman, 2002), possibly even without the need to perceive the faces consciously (Axelrod et al., 2015). As the N170 is the earliest reliable ERP index of automatic threat prioritization, it is also the most likely candidate to be an index of nonconscious emotion processing. Studies on neural correlates of nonconscious emotion processing typically use some form of “blinding technique” to prevent stimuli from being consciously perceived and to obtain a behavioral measure to test how successfully conscious perception could be prevented. The most commonly used blinding technique in studying nonconscious emotion processing is backward masking (BM; e.g., Kiss & Eimer, 2008; Pegna et al., 2011; M. L. Smith, 2012), in which the critical face stimulus is presented for a brief duration (typically between 10-30 ms). The stimulus is followed by a mask, which sometimes consists of another face with a neutral expression (e.g., Japee et al., 2009; Kiss & Eimer, 2008; Pegna et al., 2008) or a pattern mask (Doradzińska & Bola, 2024; Kiss & Eimer, 2008; M. L. Smith, 2012). The latter are often created from “scrambled” faces, i.e., small pieces of face images, randomly re-arranged to create a non-face-like random pattern of comparable luminance.

However, the results from studies using backward masked fearful faces are heterogeneous. The earliest differences between masked fearful and neutral faces have been reported between 80 and 130 ms in an magnetoencephalographic (MEG) study (Bayle et al., 2009). Some studies have reported N170 differences (Pegna et al., 2011; M. L. Smith, 2012) or differences in the same time range but over fronto-central sites (Kiss & Eimer, 2008). One study reported N170 effects for masked faces (Zotto & Pegna, 2015) but did not test whether subjects could detect these faces but whether they could discriminate between four emotional expressions. Others, however, found no N170 effects for masked fearful versus neutral faces (Qiu et al., 2022b, 2022a; Tipura et al., 2019), or, in an MEG study, an M170 only in subjects who could reliably detect the masked fearful faces (Japee et al., 2009). A recent study reported a difference between masked fearful and neutral faces that just missed the significance threshold (p = .084; Doradzińska & Bola, 2023).

Only a few studies have examined the EPN, with one study reporting significant effects (Walentowska & Wronka, 2012) and one reporting no significant effects (Doradzińska & Bola, 2023). In contrast, in studies with clearly visible faces, the EPN has more often been found to be increased for fearful compared to neutral expressions (Durston & Itier, 2021; Schindler et al., 2019; Wieser et al., 2012).

This heterogeneity is also observed in other paradigms manipulating consciousness, such as continuous flash suppression. According studies provided evidence both for (Jiang et al., 2009) and against (Schlossmacher et al., 2017) N170 modulations by nonconscious emotional expressions. Comparably, several fMRI studies presenting masked fearful and neutral faces reported positive evidence (e.g., Jiang & He, 2006; Kim et al., 2010; Liddell et al., 2005; Phillips et al., 2004; Troiani et al., 2014; Whalen et al., 1998, 2004). Others showed that these effects disappear if trial-by-trial measures of visibility are taken into account (Pessoa et al., 2006), or that they depend on target-mask interactions (Kim et al., 2010, 2016; Straube et al., 2010). Correspondingly, recent reviews have concluded that the debate about the existence of nonconscious emotion processing is not settled (Lanfranco et al., 2023; Mudrik & Deouell, 2022).

Even the seemingly simpler question of whether nonconscious faces can be differentiated from non-facial stimuli has not been unequivocally answered. One MEG study showed an enhanced M170 in response to faces compared to houses under CFS (Sterzer et al., 2009). Another CFS study presented upright and inverted faces and found the typically observed enhanced N170 for consciously perceived inverted compared to upright faces, but interestingly, it showed a reversed effect in the nonconscious condition (Suzuki & Noguchi, 2013). Two studies using inattentional blindness to prevent conscious perception either found no significant effects (Shafto & Pitts, 2015) or reported Bayesian evidence for the absence of a N170 difference between line-drawing faces and random line patterns (Dellert et al., 2021). A backward masking study found no significant N170 differences; however, the data indicated a difference that might have been overlooked due to low statistical power (N = 11; Rodríguez et al., 2012). An interesting distinction was made in a study by Harris and colleagues (2011) who presented faces, houses and blank stimuli in a sandwich masking paradigm. The authors found that the N170 did not differ between nonconscious faces and houses. However, the P1 differed between blank trials and the average across face and house trials, suggesting nonconscious low-level stimulus processing but not high-level face processing. We conclude that results regarding neural correlates of the differentiation between nonconscious facial expressions and between faces and non-faces are heterogeneous.

In addition to typical empirical problems, such as low statistical power or the use of face masks leading to interactions between target faces and masks (see Straube et al., 2010), one reason for the heterogeneity of results may be the variability in how consciousness is controlled and measured (Kerr et al., 2017; Lanfranco et al., 2023; Moors, 2019; Moors et al., 2019; Sterzer et al., 2014). The extent of nonconscious emotion processing may be overestimated in studies that measured awareness and neural correlates of threat processing in separate experimental sessions (Bayle et al., 2009; Jiang & He, 2006; Pegna et al., 2011; Troiani et al., 2014; Whalen et al., 1998), used lenient criteria to label participants as nonconscious (Kim et al., 2010, 2016; Whalen et al., 2004), or tested the absence of awareness only at the group level (Liddell et al., 2005; Phillips et al., 2004). To overcome these limitations, awareness needs to be assessed on a trial-by-trial basis with sufficiently sensitive behavioral measures. The so-called Perceptual Awareness Scale (PAS) is the most sensitive for this purpose (Sandberg et al., 2010; Sandberg & Overgaard, 2015).

The present preregistered (https://osf.io/8d3js/) ERP study was therefore designed to test nonconscious neural processing of masked fearful versus masked neutral faces with a sufficiently large sample (determined via Bayesian sampling stopping criteria) and a suited trial-by-trial assessment of subjective awareness based on the PAS. Electrophysiological analyses focused on the P1, N170, and EPN. We presented fearful faces, neutral faces, and homogenous face-shaped, monochrome control stimuli matched to each of the face’s average skin color (referred to as blanks) for 17 ms. These stimuli were followed by masks (scrambled fearful and neutral faces generated from small rectangular cutouts) after an inter-stimulus interval (ISI) of either 0 ms (ISI0) or 200 ms (ISI200). After each trial, participants were asked to respond on a 4-point PAS whether they saw nothing but the mask, saw something but nothing face-like, had a vague impression of a face, or had a clear impression of a face. For the analysis of nonconscious processing, only trials with the lowest rating were analyzed.

The primary analysis focused on fearful vs. neutral face processing under nonconscious and conscious conditions. A secondary analysis addressed nonconscious and conscious processing of neutral faces vs. control stimuli. We hypothesized that the N170 and the EPN would differentiate between conscious fearful and neutral faces but that only the N170 would show differences between nonconscious fearful and neutral faces. For the comparison between faces and blanks, we expected increased P1 and N170 amplitudes in the conscious as well as the nonconscious condition.

## 2. Methods

### Participants

We recruited a total of 68 participants at the University of Münster. Four participants were excluded due to poor EEG data (three had a signal out of range and one aborted the EEG test session) and two participants because their response bias did not meet the predefined criterion. This left a final sample of 62 participants (48 female, 13 male, 1 diverse) with a mean age of 22.97 years (*SD* = 2.46, Min = 19, Max = 31). Participants gave written informed consent and were paid or received course credit for their participation. All participants had normal or corrected-to-normal vision, were right-handed, and had no reported history of neurological or psychiatric disorders. We tested for nonconscious fearful-neutral effects using a Bayesian t-test. As registered, we tested a minimum of 40 healthy participants but stopped sampling when the Bayes factor (BF) showed more than moderate evidence in favor of a larger N170 for nonconscious fearful compared to neutral faces or in favor of a null effect (*BF*_01_ > 3 or *BF*_01_ < 1/3). We tested our stopping rule after 40 usable datasets and then after every five collected data sets (i.e., 45, 50, 55, etc.) while including participants already scheduled and removing participants with extreme response biases only after sampling was complete.

### Stimuli

As target faces, we used 40 female and 40 male individuals showing fearful and neutral expressions. This resulted in 160 different images. The target faces’ ages ranged approximately between 18 and 40 years. Based on a previous backward-masking study (Neumeister et al., 2017), the stimuli were taken from different face sets: the FACES database (Ebner et al., 2010), the NimStim set (Tottenham et al., 2009), and the Radboud Faces Database (Langner et al., 2010). Masks were created by cutting the face region of each individual image into squares of 10 × 10 pixels and rearranging these squares randomly (scramble filter of Adobe Photoshop CS6, version 13.0.1, Adobe Systems Inc., San Jose, California, USA). This resulted in 160 masks, each corresponding to one face image. In addition, artificially generated ’blank’ stimuli were created. These monochrome ovals were created by applying the soft drawing filter of Adobe Photoshop CS6 to each individual image to create control stimuli that matched the face’s average skin color. Of those 160 blank stimuli, we randomly selected 80 with the same proportions of male and female, fearful and neutral images as in the originals. The total set thus consisted of 240 target images and 160 masks. An additional set of four identities was used for practice trials.

### Procedure

The experiment was programmed and run using Matlab (version R2022a; MathWorks Inc., Natick, MA; http://www.mathworks.com) and the Psychophysics Toolbox (Version 3.0.19; Brainard, 1997; Kleiner et al., 2007). Participants underwent a backward masking procedure (see Figure 1). Each trial began with the presentation of a fixation cross in the center of the screen for a randomized duration between 500 and 700 ms. Next, the target stimulus (fearful or neutral face or blank) was presented for 17 ms. To vary the degree of conscious visibility, we manipulated the ISI between the target face and the mask: in the ISI0 condition, the mask onset coincided with the target offset, whereas in the ISI200 condition, the target offset and the mask onset were separated by a blank screen lasting 200 ms. We expected the majority of target faces to be nonconscious in the ISI0 condition and to be conscious in the ISI200 condition. We used a PAS to test this assumption. 600 ms after the offset of the target stimulus, a response prompt was shown, asking participants to rate their subjective percept on a perceptual awareness scale. Specifically, they indicated whether they noticed nothing (“nothing but the mask”; PAS = 1), saw “something but nothing face-like” (PAS = 2), had a “vague impression of a face” (PAS = 3), or a “clear impression of a face” (PAS = 4). After each key press, the selected response was surrounded by a white frame for 200 ms. Within each ISI condition, each target stimulus (fearful face, neutral face, blank) was presented twice, once with the mask generated from the corresponding neutral and fearful face, respectively. This resulted in 960 trials (640 face trials, 320 blank trials). Four different pseudorandomized trial sequences were generated by shuffling the order of trials so that the same stimulus type (face / blank), expression (fearful / neutral), and ISI (0 / 200 ms) were repeated no more than three times in a row. The four trial sequence versions were balanced across subjects. Prior to the main experiment, participants completed 32 practice trials to familiarize themselves with the stimulus presentation and the response keys.

**Figure 1:**
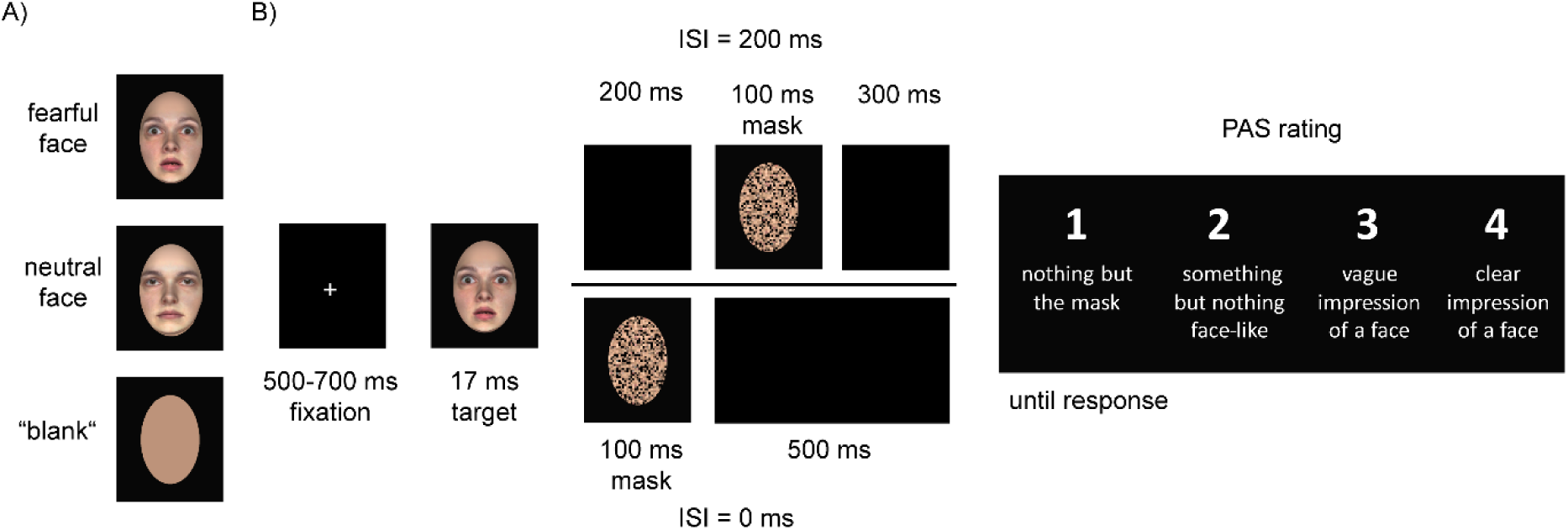
Trial sequence during the experiment. Please note that the faces shown here are not those actually used in the experiment, but were generated artificially using the FaceGen software (version 3D Print Home 2.0; Singular Inversions Inc.; https://facegen.com).

### Behavioral data analyses

Behavioral data were analyzed using Matlab (version R2022a; MathWorks Inc., Natick, MA; http://www.mathworks.com) and JASP (version 0.11.1; JASP Team, 2016). The analysis was based on each PAS level’s relative frequency and signal detection theory-based measures (SDT; Green & Swets, 1966). For the latter purpose, ratings were dichotomized, with PAS = 1 considered as “no”-responses and PAS > 1 as “yes”-responses. This two-fold approach of using a scale with four labels based on subjective perception and dichotomizing these ratings into “yes”- and “no-responses allowed us to take advantage of the optimal sensitivity of the PAS scale (Sandberg & Overgaard, 2015) while at the same time applying classical SDT measures to calculate detection sensitivity (d’) and the response criterion (c; Hautus et al., 2021). Response rates of 0 and 1 were replaced by .003125 and .996875, corresponding to 0.5 out of 160 trials (Macmillan & Kaplan, 1985). We tested participants for extremely conservative or liberal response criteria. To do so, we calculated c per participant, averaged across conditions, and then z-standardized across participants to identify extremely liberal or conservative participants, as these could lead to an under- or overestimation of nonconscious processing. All further analyses excluded participants with |z_c_| > 2.5. Next, d’ and c were analyzed using a 2 (ISI) × 2 (expression) repeated-measures ANOVA. The significance level for all F-tests was α = .05. We additionally report Bayes Factors (BF), with *BF*_01_ denoting the evidence for the null hypothesis and *BF*_10_ for the alternative hypothesis. We use the conventions from Jeffreys (1961) to interpret the results of our Bayesian analyses.

### EEG recording and preprocessing

EEG signals were recorded from 64 BioSemi active electrodes using Biosemi’s ActiView software (www.biosemi.com). Four additional electrodes measured horizontal and vertical eye movement. The recording sampling rate was 512 Hz. Offline, the data were re-referenced to average reference and band-pass filtered from 0.01 (6 dB/octave) to 40 (24 dB/octave) Hz. Recorded eye movements were corrected using BESA’s automatic eye-artifact correction method (Ille et al., 2002). Filtered data were segmented from 200 ms before stimulus onset until 1000 ms after stimulus presentation. Bad electrodes (based on visual inspection) were interpolated using a spline interpolation procedure (*M* = 2.28, *SD* = 1.36, Min = 0, Max = 6). Two participants had electrodes interpolated within the sensors of interest (one and two electrodes). Baseline correction was used at the 200 ms interval before stimulus onset.

The dependent variable for all ERP analyses were amplitudes averaged over the following time windows and channels: We preregistered to examine the mean amplitude for the P1 from 80 to 100 ms, for the N170 from 130 to 170 ms, and for the EPN from 200 to 350 ms. The preregistered sensors of interest for the P1, N170, and EPN consisted of two symmetrical occipital clusters (left P9, P7, PO7; right P10, P8, PO8). ERPs were calculated by averaging nonconscious and conscious trials separately for each target stimulus type (fearful face, neutral face, blank). Nonconscious trials were defined as trials with ISI = 0 and PAS = 1. Conscious trials were defined as trials with ISI = 200 ms and PAS > 1.

### Statistical analyses

ERPs were statistically analyzed using 2 (expression: fearful face, neutral face) by 2 (consciousness: nonconscious vs. conscious) repeated measures ANOVAs. For each ERP component, the corresponding main effects and interactions were tested. Partial eta-squared (η_P_²) was used to describe effect sizes (Cohen, 1988). Crucially, we preregistered the following planned comparisons to be tested using Bayesian t-tests: (1A) conscious fearful faces vs. conscious neutral faces; (1B) nonconscious fearful faces vs. nonconscious neutral faces. We preregistered to perform additional linear mixed-effects (LME) analysis with the amplitude of the components as dependent variables and ISI, expression, and PAS ratings as separate trial-by-trial predictors in case we obtained considerable variability of PAS ratings, which we defined as ≥ 20% of trials in the ISI0 condition with PAS ratings > 1. However, we observed on average only 5.54% of trials falling into this category and, therefore, did not perform LME analysis.

Secondary analyses were conducted to test for nonconscious face effects. Face processing can be regarded as a hierarchical process during which differential processing of facial expressions occurs at a higher level than face detection, i.e., discriminating between face and non-face stimuli (Duchaine & Yovel, 2015). Nonconscious face processing can, therefore, be understood as a “lower bar” to prove than nonconscious emotion processing. Surprisingly, even regarding nonconscious face processing, the empirical findings are mixed (Dellert et al., 2021; Navajas et al., 2013; Reiss & Hoffman, 2007; Rodríguez et al., 2012; Sterzer et al., 2009; Suzuki & Noguchi, 2013). We, therefore, performed two (stimulus type: neutral faces, blanks) by two (consciousness: nonconscious vs. conscious) repeated measures ANOVAs, restricted to the P1 and the N170. The secondary analysis was complemented by the following planned comparisons using Bayesian t-tests: (2A) conscious neutral faces vs. conscious blanks; (2B) nonconscious neutral faces vs. nonconscious blanks. Note that this secondary analysis is not statistically independent of the primary analysis; thus, the false alarm rates are also dependent.

## 3. Results

### 3.1. Behavioral analyses

#### 3.1.1 Exclusion due to extreme response biases

Two participants met the exclusion criterion of an extreme response bias, defined as |z_c_| > 2.5. They were excluded from all further analyses, leaving a sample of *N* = 62.

#### 3.1.2 Detection sensitivity d’

As predicted, we observed a significant main effect of ISI, *F*_(1,61)_ = 1513.01, *p* < .001, η_P_²= .961, characterized by a lower average d’ in the ISI0 condition (*M* = 0.59; *SD* = 0.74) compared to the ISI200 condition (*M* = 4.80; *SD* = 0.63). Contrary to our prediction that fearful faces would be detected more often than neutral faces, the main effect of expression did not reach significance, *F*_(1,61)_ = 3.31, *p* = .074, η_P_² = .051. Descriptively, d’ was higher for fearful faces (*M* = 2.73, *SD* = 2.24) than for neutral faces (*M* = 2.66, *SD* = 2.20). The interaction between ISI and expression was not significant, *F*_(1,61)_ = 0.31, *p* = .582, η_P_² = .005. Figure 2A (top) shows individual and average d’ scores.

**Figure 2:**
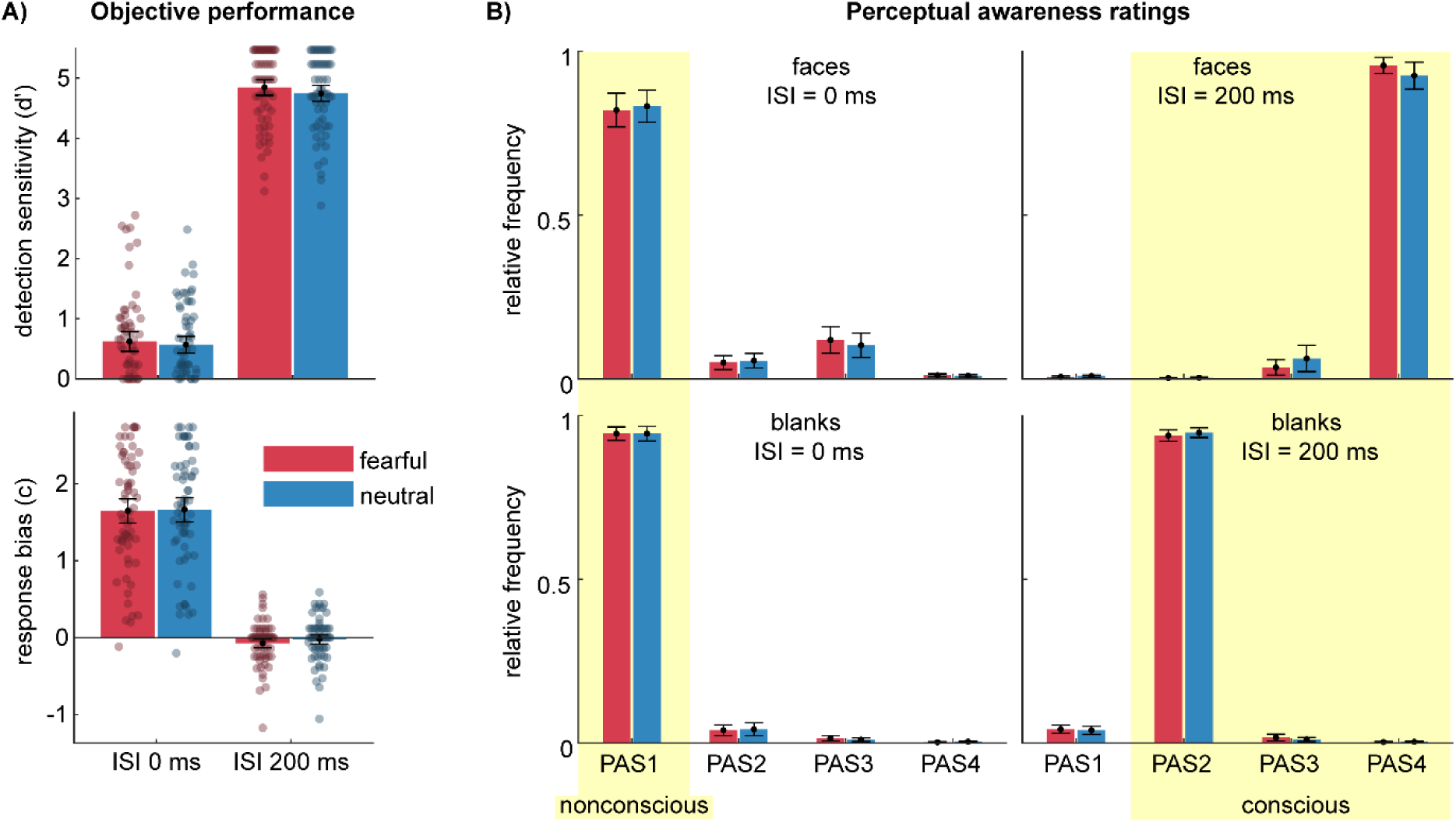
Behavioral data for fearful (red) and neutral (blue) faces and color-matched blanks. A) Objective performance measures for detection sensitivity (d’; top plot) and response bias (c; bottom plot; positive values indicate a conservative bias). Semitransparent dots represent individual participants (N = 62, i.e., after application of the exclusion criteria). B) Distribution of perceptual awareness ratings for faces (top row) and blanks (bottom row), presented with a target-mask ISI of 0 ms (left column) or 200 ms (right column). Because the blanks were constructed from individual images varying in average color and brightness, the ratings are presented separately for blanks constructed from fearful and neutral faces. The yellow rectangles highlight the conditions labeled as nonconscious and conscious for the EEG analyses. All error bars represent 95% confidence intervals of the mean.

#### 3.1.3 Response bias c

We did not have specific hypothesis regarding the response bias c. We observed a main effect of ISI, *F*_(1,61)_ = 303.69, *p* < .001, η_P_² = .833, due to more conservative response biases for ISI0 (*M* = 1.65; *SD* = 0.75) than for ISI200 (*M* = - 0.05; *SD* = 0.29). The main effect of expression was not significant, *F*_(1,61)_ = 3.11, *p* = .083, η_P_² = .049, nor was the interaction of expression and ISI significant, *F*_(1,61)_ = 1.04, *p* = .312, partial η_P_² = .017. Figure 2A (bottom) illustrates individual and average response bias scores.

#### 3.1.4 PAS ratings

Figure 2B shows the distribution of PAS ratings to illustrate which conditions were labeled as nonconscious or conscious for the EEG analyses. We did not perform any statistical analyses of these distributions.

### 3.2. ERPs

#### 3.2.1. P1

For the P1, we observed no main effects of expression (*F*_(1,61)_ = 2.45, *p =* .123, η_P_² = .039, see Figure 3) or consciousness (*F*_(1,61)_ = 2.20, *p =* .144, η_P_² = .035), nor an interaction between expression and consciousness (*F*_(1,61)_ = 0.03, *p =* .869, η_P_² < .001). The planned Bayesian t-test per consciousness condition showed moderate evidence against a fearful-neutral difference for nonconscious trials (*BF*_01_ = 4.048, error < 0.001) and moderate evidence against a fearful-neutral difference for conscious trials (BF_01_ = 3.390, error < 0.001). In secondary analyses of nonconscious face effects, we observed a main effect of stimulus type (*F*_(1,61)_ = 27.33, *p <* .001, η_P_² = .309, see Figure 4), but no significant effect of consciousness (*F*_(1,61)_ = 3.86, *p =* .054, η_P_² = .059). There was an interaction between stimulus type and consciousness (*F*_(1,61)_ = 13.95, *p <* .001, η_P_² = .186) due to a stronger face-blank difference in the conscious condition. The planned Bayesian t-test per consciousness condition revealed that there was anecdotal evidence against a face-blank difference for nonconscious trials (*BF*_01_ = 1.608, error < 0.001) but extreme evidence for a difference in conscious trials (*BF*_10_ = 203,430.114, error < 0.001).

**Figure 3.**
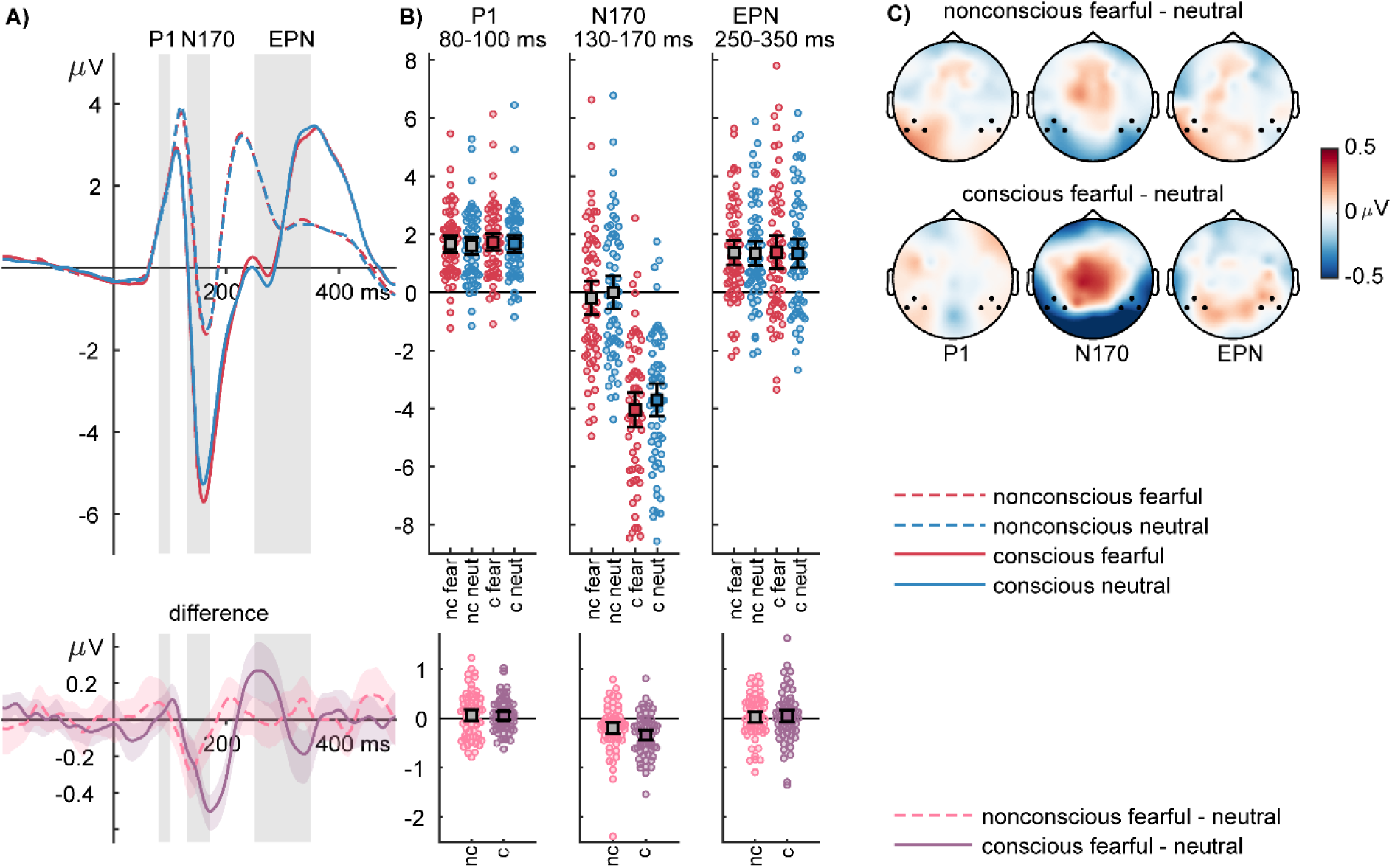
P1, N170, and EPN expression effects. **A)** Top panel: ERP waveforms showing the time course for fearful (red) and neutral (blue) faces in the nonconscious (ISI0, PAS = 1; dashed lines) and the conscious (ISI200, PAS > 2; solid lines) condition. Bottom panel: difference waves (fearful – neutral) for the nonconscious (pink) and conscious (purple) conditions. Shaded areas indicate the 95% bootstrap confidence interval for the average difference. Gray areas mark the intervals of interest used to compute average amplitudes and topographies. **B)** Individual amplitudes and error bar plots with 95% confidence intervals for each interval of interest, averaged over the channels of interest highlighted in the topography plots). **C)** Scalp topographies depict the amplitude difference between fearful and neutral expressions for nonconscious (top row) and conscious (bottom row) conditions.

**Figure 4.**
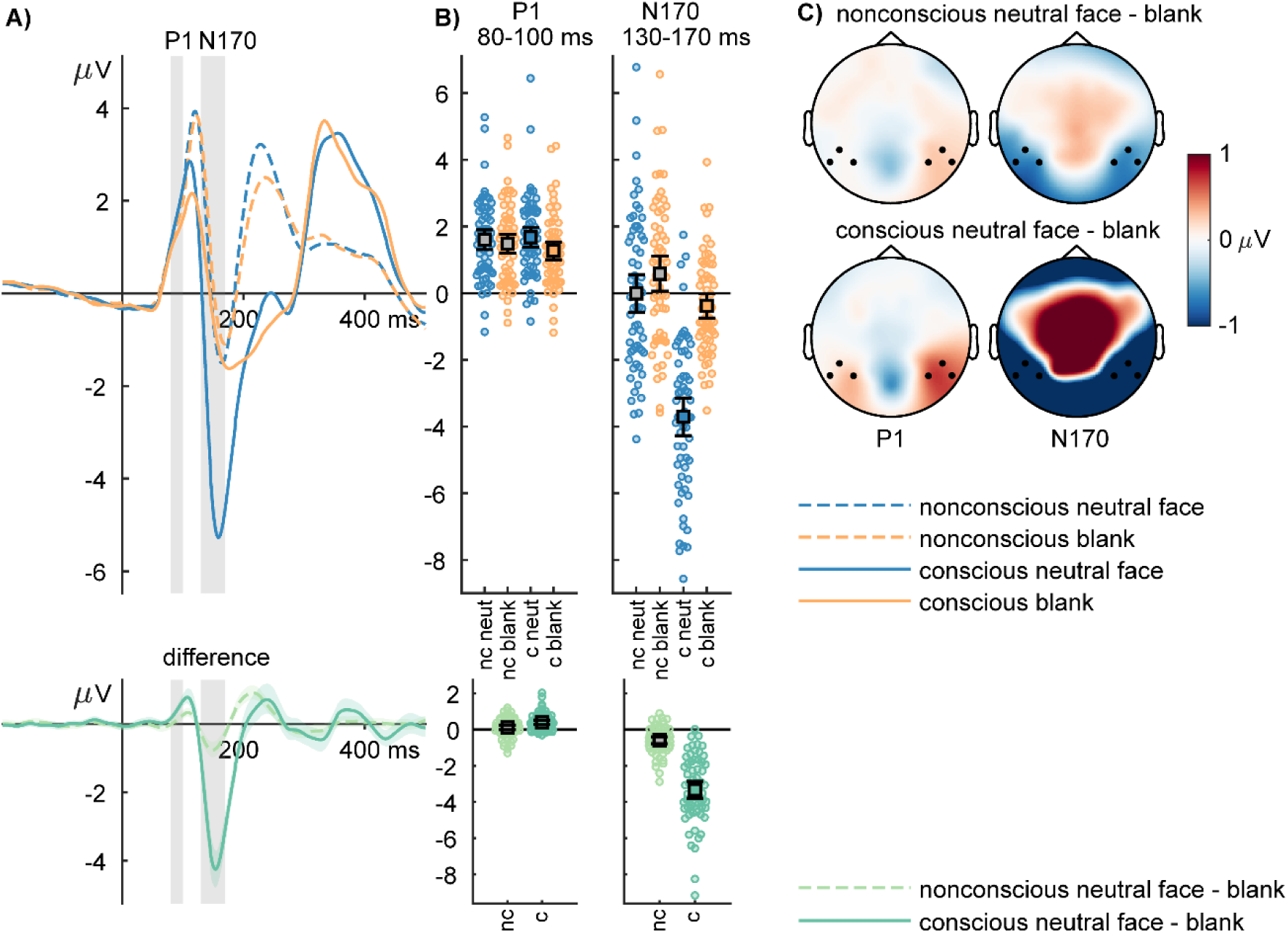
P1 and N170 face-blank effects. **A)** Top panel: ERP waveforms showing the time course for neutral faces (blue) and “blanks” (yellow) in the nonconscious (ISI0, PAS=1; dashed lines) and the conscious (ISI200, PAS>2; solid lines) condition. Bottom panel: difference waves (neutral face - blank) for the nonconscious (light green) and conscious (darker green) conditions. Shaded areas indicate the 95% bootstrap confidence interval for the average difference. Gray areas mark the intervals of interest used to compute average amplitudes and topographies. **B)** Individual amplitudes and error bar plots with 95% confidence intervals for each interval of interest, averaged across the channels of interest highlighted in the topography plots). **C)** Scalp topographies showing the amplitude difference between neutral faces and blanks for nonconscious (top row) and conscious (bottom row) conditions.

##### Exploratory analyses of mean P1 amplitudes around the peak (105 to 125 ms)

Because the collapsed P1 amplitude peak did not fall within the preregistered time window, we performed additional, unregistered analyses with a symmetric time window around the P1 peak (115 ms ± 10 ms). We still observed no main effects of expression (*F*(1,61) = 0.89, *p* = .350, η_P_² = .014) and no interaction between expression and consciousness (*F*(1,61) = 2.62, *p* = .111, η_P_² = .041). Although we found an effect of consciousness (*F*(1,61) = 52.39, *p* < .001, η_P_² = .462), with larger P1 amplitudes in the nonconscious condition, we consider this effect uninterpretable due to the different ISIs underlying conscious and nonconscious trials (see Figure 5A). Bayesian t-tests, analogous to the tests performed in the preregistered interval, revealed anecdotal evidence for the absence of a fearful-neutral difference among nonconscious faces, *BF*_01_ = 2.184, error < 0.001, and moderate evidence for the absence of a fearful-neutral difference among conscious faces, *BF*_01_ = 6.510, error < 0.001.

**Figure 5.**
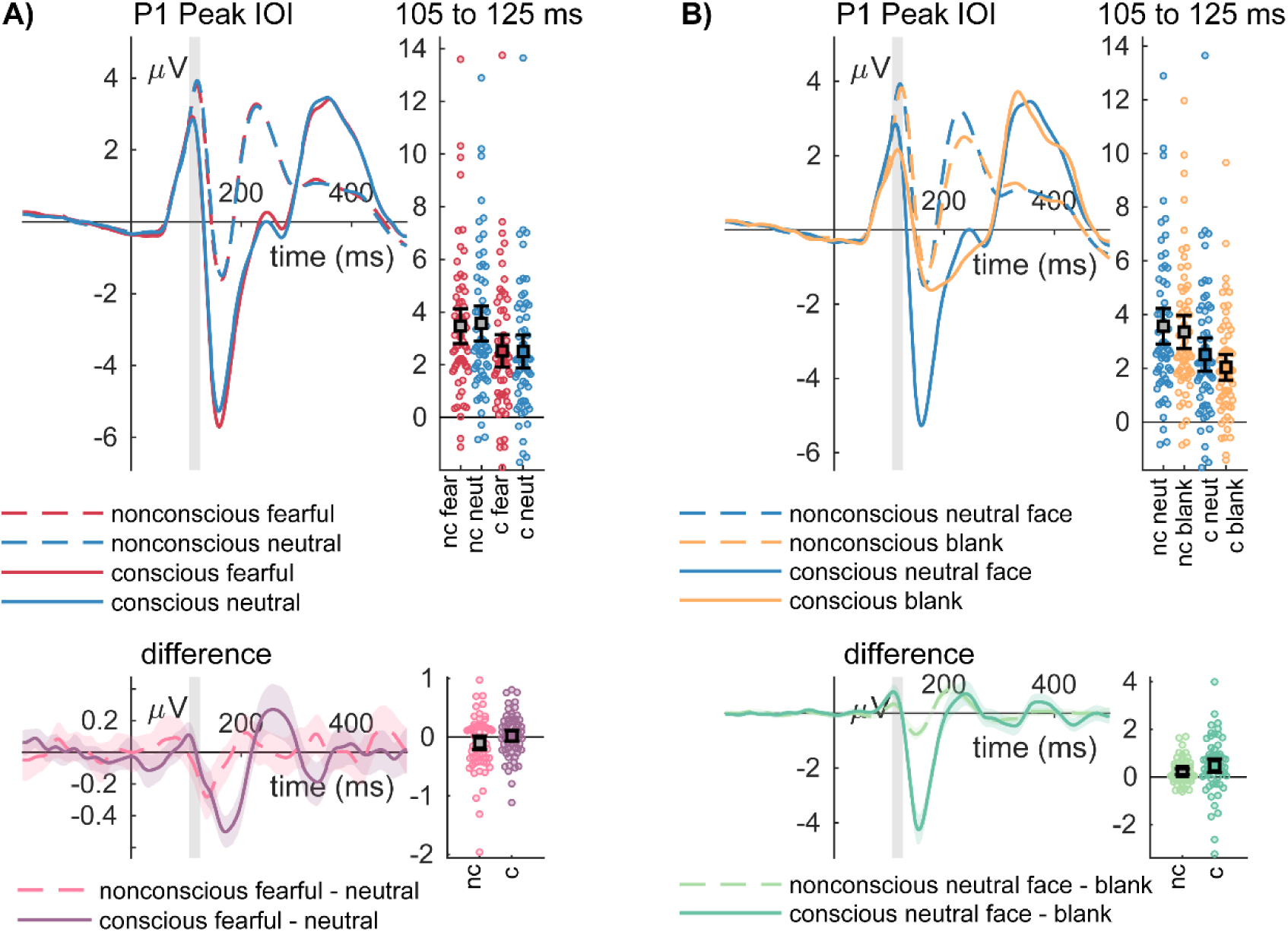
P1 emotion and expression effects centred on the peak of the grand-average P1. **A)** Top panel: ERP waveforms showing the time course for fearful (red) and neutral (blue) faces in the nonconscious (ISI0, PAS=1; dashed lines) and the conscious (ISI200, PAS>2; solid lines) condition. Bottom panel: difference waves (fearful – neutral) for the nonconscious (pink) and conscious (purple) conditions. **B)** Top panel: ERP waveforms show the time course for neutral faces (blue) and “blanks” (yellow) in the nonconscious (ISI0, PAS=1; dashed lines) and the conscious (ISI200, PAS>2; solid lines) condition. Bottom panel: difference waves (neutral face - blank) for the nonconscious (light green) and conscious (darker green) conditions. Shaded areas indicate the 95% bootstrap confidence interval for the average difference. Gray areas mark the intervals of interest.

In secondary analyses of nonconscious face effects, we observed main effects of stimulus type (*F*_(1,61)_ = 14.25, *p <* .001, η_P_² = .189) and consciousness (*F*_(1,61)_ = 63.51, *p <* .001, η_P_² = .509). The interaction between stimulus type and consciousness did not reach significance (*F*_(1,61)_ = 3.41, *p =* .069, η_P_² = .053; see Figure 5B). Bayesian t-tests, analogous to the tests performed in the preregistered interval, revealed strong evidence for a difference between nonconscious neutral faces and nonconscious blanks, *BF*_10_ = 26.760, error < 0.001, and between conscious neutral faces and conscious blanks, *BF*_10_ = 12.369, error < 0.001.

#### 3.2.2. N170

For the N170, we observed main effects of expression (*F*_(1,61)_ = 39.46, *p <* .001, η_P_² = .393) and consciousness (*F*_(1,61)_ = 315.38, *p <* .001, η_P_² = .838). The interaction between expression and consciousness did not reach significance (*F*_(1,61)_ = 3.29, *p =* .075, η_P_² = .051). The planned Bayesian t-test per consciousness condition showed strong evidence for a fearful-neutral difference for nonconscious trials (*BF*_10_ = 12.64, error < 0.001) and extreme evidence for a fearful-neutral difference for conscious trials (*BF*_10_ = 233,431.344, error < 0.001). Note that the Bayes factor for nonconscious fearful-neutral differences is higher than the one used for the sampling stopping rule (*BF* > 3) because the stopping rule was computed before excluding two participants based on their response criteria.

In secondary analyses of nonconscious face effects, we observed main effects of stimulus type (*F*_(1,61)_ = 178.63, *p <* .001, η_P_² = .745, see Figure 4) and consciousness (*F*_(1,61)_ = 178.63, *p <* .001, η_P_² = .732). There was an interaction between stimulus type and consciousness (*F*_(1,61)_ = 159.96, *p <* .001, η_P_² = .724) due to a stronger face-blank difference in the conscious condition. The planned Bayesian t-test per consciousness condition revealed extreme evidence for a face-blank difference for nonconscious trials (*BF*_10_ = 1.511×10^6^, error < 0.001) and conscious trials (*BF*_10_ = 3.538×10^17^, error < 0.001), respectively.

Additionally to these preregistered analyses, we tested whether the observed N170 differences between nonconscious fearful and neutral expressions and between neutral faces and blanks correlate with the individual response bias c. No significant correlation was observed (fear - neutral vs. c: r = .0998, p = .440; neutral – blank vs. c: r = −.1532, p = .234).

#### 3.2.3. EPN

For the EPN, we observed no main effect of expression (*F*_(1,61)_ = 1.34, *p =* .251, η_P_² = .022), but an effect of consciousness (*F*_(1,61)_ = 40.70, *p <* .001, η_P_² = .400). There was no interaction between expression and consciousness (*F*_(1,61)_ = 0.01, *p =* .932, η_P_² < .001). The planned Bayesian t-test per consciousness condition showed moderate evidence against fearful-neutral differences for nonconscious trials (*BF*_01_ = 5.405, error < 0.001) and conscious trials (*BF*_01_ = 5.181, error < 0.001), respectively.

## 4. Discussion

The present ERP study was designed to test the nonconscious processing of masked faces with fearful versus neutral expressions. To obtain a sufficiently large sample, we determined the stopping rule based on Bayesian evidence for or against nonconscious expression effects in the N170 component. The PAS was used to ensure accurate trial-by-trial assessment of subjective awareness. We observed strong evidence for nonconscious differential processing of fearful versus neutral faces in the N170 but not in other ERP components (P1, EPN). Furthermore, we found extreme evidence for nonconscious face processing (neutral faces vs. blank stimuli), even though participants subjectively reported experiencing “nothing but the mask”.

Our results show that with high statistical power significant face- and expression-specific effects on the N170 can be observed even when participants have no subjective experience of a stimulus other than the mask. We did not find an influence of the response criterion, suggesting that the N170 effects are not produced by those individuals who have a highly conservative criterion (i.e., responding with “nothing but the mask” even when perceiving a glimpse of a stimulus).

Some previous backward masking studies have found evidence for N170 (or M170) emotion effects (e.g., Pegna et al., 2011; M. L. Smith, 2012; Zotto & Pegna, 2015), while others have not (Japee et al., 2009; Qiu et al., 2022a, 2022b, 2023), or report a statistical trend (Doradzińska & Bola, 2023). Null findings are difficult to interpret and may depend on statistical power. The results of Japee and colleagues (2009) indicate a small but statistically insignificant increase of the M170 for nonconscious fearful faces. However, this test was based on only 12 participants, suggesting that potential effects may have been overlooked. The studies by Qiu and colleagues (2022a, 2022b, 2023) are comparably high-powered (22 ≤ N ≤ 42) but were not specifically designed to test N170 effects. Instead, they examined the effects of shifting spatial attention to nonconscious fearful faces and thus presented faces lateralized. Lateralized faces, however, even if presented unmasked, show only attenuated or no N170 enhancement by fearful compared to neutral faces (Schindler, Meyer, et al., 2022). Thus, these studies may not have provided the optimal testing ground to examine N170 effects for masked fearful faces. Our results provide the first strong Bayesian evidence for a nonconscious fearful-neutral differentiation in the N170 component. This observation is consistent with more general findings that such fearful-neutral differences are found in the N170 independent of various experimental manipulations, as described in the introduction (e.g. Bruchmann et al., 2020; Mühlberger et al., 2009; Schindler, Caldarone, et al., 2020; Schindler et al., 2019; E. Smith et al., 2013; Walentowska & Wronka, 2012; Wieser et al., 2012). However, while feature-based attention and attentional load only have a limited effect on fearful-neutral N170 differences, the awareness manipulation in the current study has a strong impact as indexed by a 1.73-fold increase in the average fearful-neutral difference under conscious compared to nonconscious conditions. This factor is a very conservative estimate as it only pertains to the averaged interval of interest. A comparison of the fearful-neutral difference waves in Figure 3A shows that nonconscious differences are not only considerably smaller but appear to be shorter in duration. Specifically, the differences are observed on the descending, early branch of the N170, peaking at about 140 ms, while the N170 itself peaks at about 165 ms. Interestingly, the N170 has been shown to be modulated by awareness of faces (Dellert et al., 2021; Roth-Paysen et al., 2022; Shafto & Pitts, 2015), leading to the speculation that only the early phase of the N170 may index nonconscious processing while consciousness emerges around the time of the peak of the N170.

For the comparison between faces and control stimuli, we also found evidence for strong effects during the N170 processing stage, suggesting that at least the clear physical differences between neutral faces and blanks are processed without subjective awareness at this early stage. However, stimulus awareness enhanced this difference (by a factor of 5.6 in the interval of interest) in accordance with the involvement of the N170 in conscious awareness of faces (Dellert et al., 2021; Roth-Paysen et al., 2022; Shafto & Pitts, 2015). Taken together, the results for the N170 suggest that only some processing is partially preserved under masking conditions, while awareness potentiates both emotion and face effects.

In contrast to the N170 results and in line with our preregistered hypotheses, we observed no differences between fearful and neutral faces in the P1, in either conscious or nonconscious trials. The prediction of null effects was based on the mixed findings in studies with clearly visible faces (Itier & Taylor, 2004; Neumann et al., 2011; but see Ganis et al., 2012; Schindler, Tirloni, et al., 2021) and expressions (for a review see Schindler & Bublatzky, 2020). We assumed that very high statistical power is needed to reliably detect P1 increases for fearful faces, even for clearly visible stimuli. However, even experiments with sufficient power to detect P1 differences between conscious fearful and neutral faces may still fail to show effects in nonconscious conditions, assuming that nonconscious effects are again smaller than conscious effects. Consistent with this assumption, the exploratory analysis of nonconscious faces vs. blanks centered on the P1 peak revealed a significant difference that was again about half as large as the corresponding difference in the conscious condition. Faces and blanks differ strongly in low-level visual information, which is likely to be processed even more automatically than configural facial expressions associated with the N170. Possible P1 differences between emotional expressions can at least partly be explained by their differences in low-level properties (Bruchmann et al., 2020; Schindler, Wolf, et al., 2021). Further studies comparing facial and non-facial stimuli with better control of low-level information are needed to clarify the contributions of the P1 and N170 stages to nonconscious processing.

Finally, we found no effects of the facial expression on the EPN, in either conscious or nonconscious trials. Regarding nonconscious processing, only a few studies have examined the EPN, with one study reporting effects (Walentowska & Wronka, 2012). In contrast, studies with clearly visible faces have more often found that the EPN is increased for fearful compared to neutral expressions (Durston & Itier, 2021; Schindler et al., 2019; Wieser et al., 2012). However, the differentiation between fearful and neutral faces depends on sufficient feature-based and spatial attention and exposure times (Schindler, Bruchmann, et al., 2020; Schindler, Richter, et al., 2022; Steinweg et al., 2021; Vormbrock et al., 2023). It is possible that the very short exposure (17 ms) led to insufficient processing of relevant face information, even in trials where participants reported a clear impression of a face. While the PAS ratings clarifiy the conscious perception of facial information, we cannot conclude whether more relevant information was processed during this stage to differentiate between expressions, i.e., whether a reported “clear perception of a face” also indicates the clear perception of the expression of that face. Based on original ideas about early attentional selection processes (Schupp et al., 2004), a previous study showed a temporal sensitivity of the EPN toward a preceding demanding task (Schindler, Caldarone, et al., 2020). The authors suggested that the EPN indexes a ‘bottleneck’ of further emotion processing that is constrained by concurrent resource-dependent tasks. Future studies with additional trial-wise reports of the degree of emotional expression processing are needed to draw firm conclusions about the absence of EPN effects.

## Conclusion and outlook

This study showed that fearful faces can be differentiated from neutral faces and faces from non-facial control stimuli at the level of the N170 without subjective awareness of stimulus presentation. While our study suggests that expression and facial information can be neurally processed when backward masking is used to prevent awareness, several questions remain to be answered. Even a high-powered study needs to be replicated with different stimuli and samples. Furthermore, if replicated, it remains to be seen whether similar results will be obtained when using inherently neutral faces associated with emotional relevance (e.g., fear-conditioned faces). In addition, future studies should investigate the contributions of low-level physical differences and higher-level configural differences to nonconscious P1 and N170 effects based on studies with sufficient power to investigate early ERP components. Finally, future studies need to clarify the precise conditions for eliciting the EPN, such as the role of emotional expression perception and stimulus timings.

